# FUS Facilitates Gene Body Export from Nuclear Condensates

**DOI:** 10.1101/2023.05.25.542265

**Authors:** Seungha Alisa Lee, Emily Gini Uh, Sammantha Sae-Young Kim, Michael Zelko, Hojoong Kwak

**Affiliations:** Department of Molecular Biology and Genetics, Cornell University, Ithaca NY, 14850

## Abstract

Membraneless nuclear condensates are an emerging model of transcriptional regulation that involves spatially clustered enhancers and genes forming transcription condensates (TCs). We used a physical isolation approach and DNA/nascent RNA analysis to investigate the global organization of the genome in TCs. Comparative analysis of the two revealed the dynamic nature of the genome, with stable localization of promoters, dynamic recruitment of enhancers, and dynamic export of gene bodies out of TCs upon gene activation. Our findings also suggest that the RNA binding protein FUS undergoes decondensation upon nascent RNA binding, facilitating gene body export out of TCs.

## INTRODUCTION

FUS (Fused in Sarcoma) is an RNA binding protein that was originally discovered as an oncogene following the identification of a chromosomal translocation in myxoid liposarcoma (Rabbitts et al. 1993). In normal cells, FUS regulates RNA transcription and processing. It binds to RNA polymerase II (Pol II), general transcription factors (TFs) and specific TFs (Schwartz et al. 2015). In addition to its involvement in transcription, FUS plays a critical role in RNA processing and splicing (Yu et al. 2015). Another important function of FUS is its involvement in the transport of mRNA between the cytoplasm and the nucleus, such as the transfer of mRNA between neuronal dendrites and dendritic spines. This function is essential for neuronal cell maturation, plasticity, and maintenance of dendrite integrity.

FUS has also been implicated in several neurodegenerative diseases, including ALS and frontotemporal dementia, where it has been observed to form cytoplasmic aggregates (Schwartz et al. 2015; Maharana et al. 2018; Qamar et al. 2018). However, within the nucleus, FUS is less prone to condensation due to the RNA-rich environment (Maharana et al. 2018; Qamar et al. 2018; Chatterjee et al. 2022). RNA molecules act as buffers, preventing pathological phase separation of FUS in the nucleus (Maharana et al. 2018). Upon inhibition of global transcription, FUS redistributes to subnuclear condensate domains (An et al. 2019). The specific population of nuclear RNA responsible for this behavior remains unclear, as does whether FUS continues to co-localize with other condensates in the nucleoplasm. Regardless, these findings suggest that FUS may regulate gene expression through its condensation-decondensation dynamics.

Phase condensation in the nucleus has recently been highlighted as a novel model of gene expression regulation in which low-affinity, multivalent interactions play a critical role (Hyman et al. 2014; Banani et al. 2017; Hnisz et al. 2017; Shin and Brangwynne 2017; Langdon et al. 2018; Sharp et al. 2022). This concept, known as transcription condensates (TCs), is supported by microscopic and biophysical evidence showing that highly active transcriptional compartments are formed by spatially clustered enhancers or super-enhancers (Boehning et al. 2018; Cho et al. 2018; Lu et al. 2018). These condensates contain DNA and RNA binding factors, enzymes, and substrates that facilitate cooperative gene activation. Single-molecule RNA imaging has detected transcriptional bursts in super-enhancers within phase condensates of protein factors and nuclear RNAs (Fukaya et al. 2016; Chong et al. 2018; Sabari et al. 2018; Li and Pertsinidis 2021). In vitro studies have successfully reconstituted these structures using key transcription and chromatin factors with intrinsically disordered regions (IDRs) (Wright and Dyson 2015; Hnisz et al. 2017; Sabari et al. 2018; Cho et al. 2018; Baudement et al. 2018).

Understanding the factors that regulate chromatin within TCs is a fundamental question in the field. While disordered proteins are considered the major players, nucleic acid polymers, such as RNA molecules, have emerged as potential drivers of phase condensation (Andersson and Kedersha 2009; Van Treeck et al. 2018; Garcia Jove et al. 2019). Protein-RNA interactions, as observed in cytoplasmic RNA granules, are also critical in nuclear condensates (Soutourina et al. 2011; Lu et al. 2018; Boehning et al. 2018; Tian et al. 2020; Roden and Gladfelter 2021; Chen and Mayr 2022). Such interactions involving RNA-binding proteins may potentially affect transcription by regulating condensation, although the precise effects on TCs remain to be fully understood.

Given the condensation properties of FUS, a known RNA-binding protein, and the role of transcription condensates (TCs) in RNA regulation, it’s conceivable that FUS could act as a regulator of TCs. The well-documented functions of FUS in transcription and RNA processing, complemented by extensive research on its phase condensation potential, lend credence to this potential regulatory association with TCs. Despite this evidence, significant gaps remain in our understanding of TCs and FUS within the nucleus. These include questions regarding the specific genes and super-enhancers under transcriptional regulation within TCs, the dynamics of chromatin and transcription within TCs, and the influence of FUS condensation-decondensation dynamics on these operations.

This lack of understanding is a significant barrier to identifying TC-mediated regulation and expanding our understanding of chromatin dynamics, both within and beyond condensates (Bulger and Groudine 2011; Hnisz et al. 2013; Whyte et al. 2013; Hnisz et al. 2017). While methods such as salt-induced condensation have been tested (Baudement et al. 2018), the field currently lacks a standardized protocol for defining, isolating, or enriching TCs for comprehensive genomic analysis. This deficiency poses a significant challenge to progress in understanding these important regulatory mechanisms.

To address these questions, we developed a physical isolation strategy for mapping chromatin within nuclear TCs, inspired by techniques used to map RNA in cytoplasmic condensates. Using this approach, we analyzed the DNA and nascent RNA components of TCs genome-wide and identified genes, enhancers, and non-coding RNAs enriched in TCs. Through comparative analysis of these components, we revealed the dynamic properties of TCs during gene activation. Additionally, we discovered that FUS-nascent RNA condensation is mutually buffered, and that FUS facilitates the redistribution of elongating RNA polymerase II (Pol II) complexes out of TCs.

## RESULTS & DISCUSSION

We isolated nuclear condensates by performing light nuclease digestion on the nuclei and sedimenting a fraction that should contain TC and other nuclear condensates in HeLa cells (Fig. 1A). We optimized the Micrococcal nuclease (MNase) digestion conditions to maximize the DNA size difference between the pellet and the soluble chromatin (Fig. S1). The pelleted nuclear condensate fraction was resistant to digestion, while the DNA in the soluble fraction was digested into smaller fragments. The condensate fraction (NC) was enriched with the active histone mark acetylated histone 4 (H4ac), which was approximately 4-8 fold higher than in the whole nuclei (Fig. 1B). This suggests that the isolated nuclear condensates are predominantly active TC.

**Fig 1.**
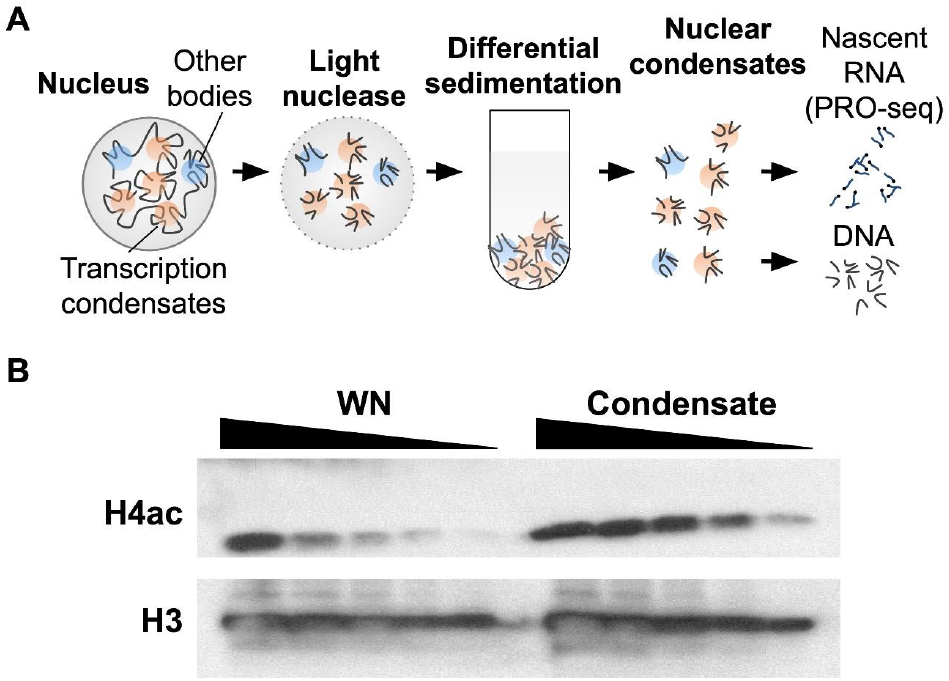
Physical isolation of transcription condensates (TC) in HeLa cells. **A**. Schematics of physical isolation. **B**. Enrichment of active histone marks in the nuclear condensate (NC) fraction. Western blots of acetylated histone 4 (H4ac) in decreasing amount (2 fold dilutions) of input whole nuclei (WN) and the condensate fraction derived from equivalent nuclei count. Histone 3 (H3) serves as a loading control.

We isolated and analyzed the DNA and nascent RNA of TC using DNA-seq and PRO-seq (Kwak et al. 2013) (Fig 1A). DNA profiling enabled us to identify the enrichment of all chromatin in TC, while nascent RNA analysis allowed us to specifically identify the actively transcribed chromatin of TC (Fig S2). To confirm the presence of active RNA polymerases in our isolated TC fractions, we performed a biotin-nucleotide run-on assay, which demonstrated compatibility with PRO-seq (Fig S3). Our PRO-seq analysis of both whole nuclei and TC revealed specific loci, including the MYC oncogene, lncRNAs, and super-enhancers, that were enriched in TC (Fig S4). Moreover, the TC fraction showed high enrichment of actively transcribed histone genes, which is consistent with previous studies on the localization of active histone genes in nuclear Histone Locus Bodies (Nizami et al. 2010) (Fig S5).

Next, we conducted a genome-wide analysis of TC DNA-seq and TC PRO-seq using ChromHMM regions to examine the distribution of genomic elements (Fig. 2, Fig. S6). As expected, both DNA-seq and PRO-seq analyses revealed a significant enrichment of promoters and enhancers within the TC (Fig. 2A: Prm, Enh). Consistent with the high enrichment of H4ac, heterochromatin DNA (Het) was depleted in the TC. Although the DNA in the transcribed gene body regions (Txn) was enriched in TC (DNA-seq), the nascent RNAs of the Txn region were not enriched (PRO-seq). Furthermore, the average metagene profile of scaled gene bodies in PRO-seq data revealed a depletion of the gene body in the TC relative to the whole nuclei (Fig. 2B). These results suggest that although gene body regions primarily reside in the TC, the actively transcribed population of the gene bodies is located outside of the TC.

**Fig 2.**
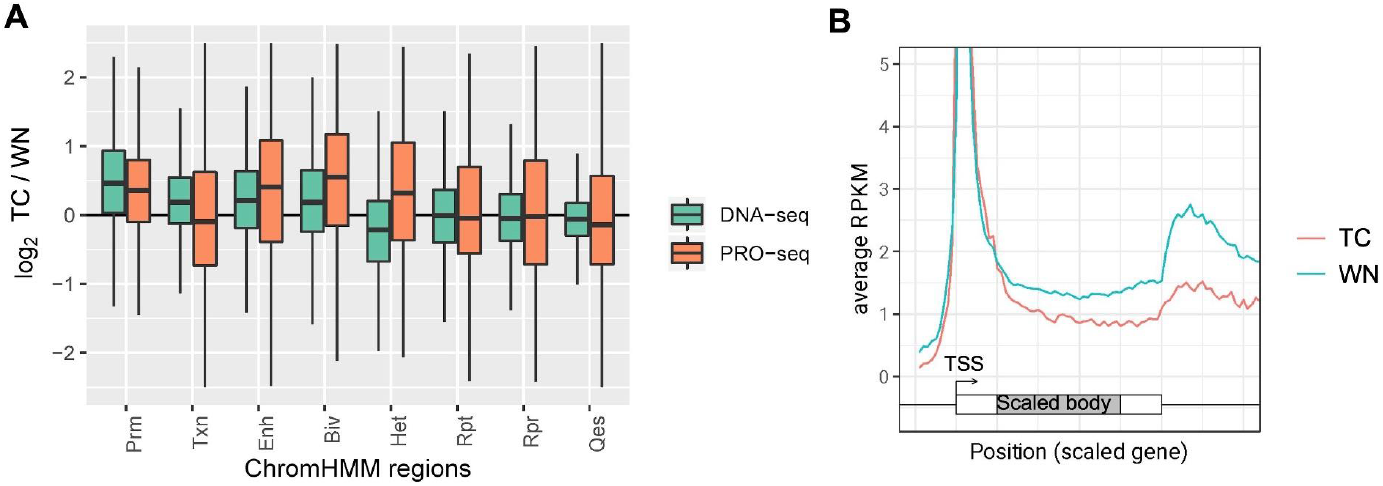
Global landscape of chromatin enrichment in TC. **A**. Distribution of the ratio between TC and whole nuclei (WN) of ChromHMM regions in DNA-seq and PRO-seq. DNA-seq measures all chromatin in the fractions, and PRO-seq measures active chromatin in the fractions (see Fig S6). ChromHMM regions are promoters (Prm), transcribed gene bodies (Txn), enhancers (Enh), bivalent chromatin (Biv), heterochromatin (Het), repetitive region (Rpt), repressed chromatin (Rpr), and quiescent chromatin (Qes). **B**. Average metagene profile of PRO-seq data depicting a scaled gene body in TC and WN. The first and the last 500 bp of the genes are not scaled (white box), while the remaining mid-gene bodies are scaled (gray box).

We propose a hypothesis that gene bodies are dynamically exported outside of the TC upon transcription activation. To investigate the distribution of genomic elements, we conducted a genome-wide analysis of TC DNA-seq and TC PRO-seq using ChromHMM regions (Ernst and Kellis, 2012) (Fig 2). Both DNA-seq and PRO-seq analyses revealed a significant enrichment of promoters and enhancers within the TC (Fig 2A: Prm, Enh). The TC exhibited a depletion of heterochromatin DNA (Het), consistent with the high enrichment of H4ac. Although DNA in the transcribed gene body regions (Txn) was enriched in TC (DNA-seq), the nascent RNAs of the Txn region were not enriched (PRO-seq). Moreover, the average metagene profile of scaled gene bodies in PRO-seq data demonstrated a depletion of the gene body in the TC relative to the whole nuclei (Fig 2B). These results suggest that although gene body regions primarily reside in the TC, the actively transcribed population of the gene bodies is located outside of the TC. Hence, we propose that gene bodies are dynamically exported outside of the TC upon transcription activation.

Closer examination of enhancers and other regions through comparative analysis of DNA-seq and PRO-seq revealed their dynamics. Although enhancer DNA is already enriched in the TC, it shows even greater enrichment in PRO-seq (Fig 2A, Fig S6), suggesting that enhancers transcribing eRNAs dynamically move into the TC upon activation. This observation led us to hypothesize that enhancers move into TCs upon activation, as eRNAs are a hallmark of enhancer activity. Bivalent promoters (H3K4me3 and H3K27me3) exhibit similar patterns to enhancers (Fig 2A; Biv) and suggest dynamic movement into TC upon activation. Although heterochromatin regions are transcriptionally repressed, they are known to contain regions that can become transcriptionally activated. These activatable heterochromatin regions may also be dynamically recruited into the TC, as PRO-seq data shows higher enrichment than DNA-seq data in TC (Fig 2A; Het).

We investigated the dynamics of gene body chromatin movement in and out of TC during transcription, suggesting that TCs are specialized for enhancer activation and transcription initiation. However, the transcribed RNA eventually needs to be exported from the compartment and the nucleus. We hypothesized that protein-RNA interactions drive gene body export from TC. FUS, an RNA-binding protein known to form phase condensates autonomously and regulated by RNA molecules, was examined to understand its role in gene body export. To test our hypothesis, we treated cells with Flavopiridol (FP), a pharmacological inhibitor of Pol II elongation (Chao and Price, 2001), to deplete nascent RNA. FP treatment triggered the rapid formation of nuclear phase condensates by FUS within 30 minutes (Fig 3A, Supplementary Media). Interestingly, the gene body regions became relatively more enriched in the TC fraction upon FP treatment (Fig S7), consistent with our model that transcription elongation drives the gene body out of TC. Additionally, we observed an increased amount of FUS in the TC fraction after FP treatment (Fig 3B). These findings confirm our previous model and further indicate that nascent RNA in the nucleus buffers FUS from phase condensation.

**Fig 3.**
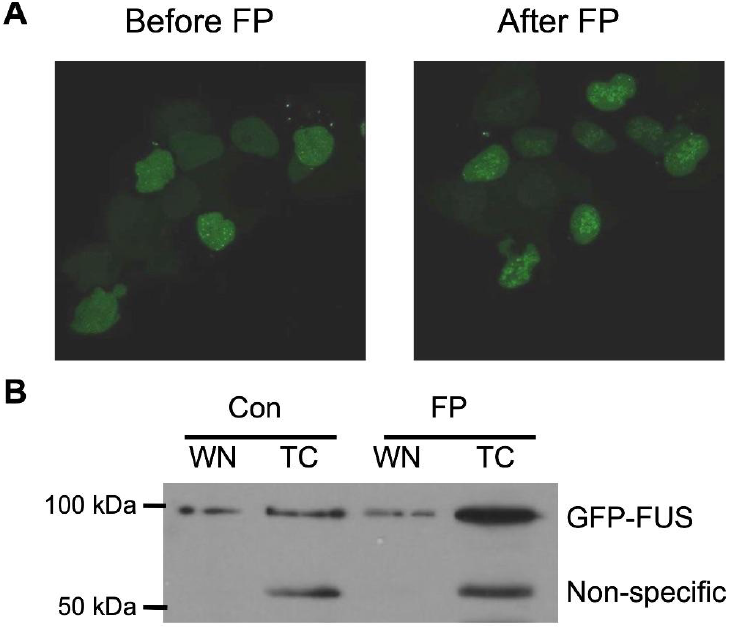
FUS condensation upon nascent RNA depletion in the nucleus. **A**. Fluorescence microscope of GFP-FUS expressing HEK293 cells before and after flavopiridol (FP) treatment. **B**. Immunoblots showing the enrichment of GFP-FUS (102 kDa) in TC after FP treatment and control (Con). Non-specific bands serve as loading controls.

If nascent RNA prevents FUS condensation, FUS may also have a role in preventing nascent RNA from condensing while assisting its movement out of nuclear condensates. Therefore, we proposed that FUS may assist in exporting nascent RNA and related gene body chromatin out of TC during the elongation process. To test this, we compared TC PRO-seq data between HEK293 cells overexpressing FUS and wild-type HEK293 cells (Fig 4A). Both FUS-overexpressing and wild-type cells displayed similar gene body patterns in the whole nuclei PRO-seq. However, FUS-overexpressing cells exhibited a reduced gene body profile compared to the wild-type cells when examining the PRO-seq data within TC, supporting our hypothesis that FUS plays a role in exporting the gene body.

**Fig 4.**
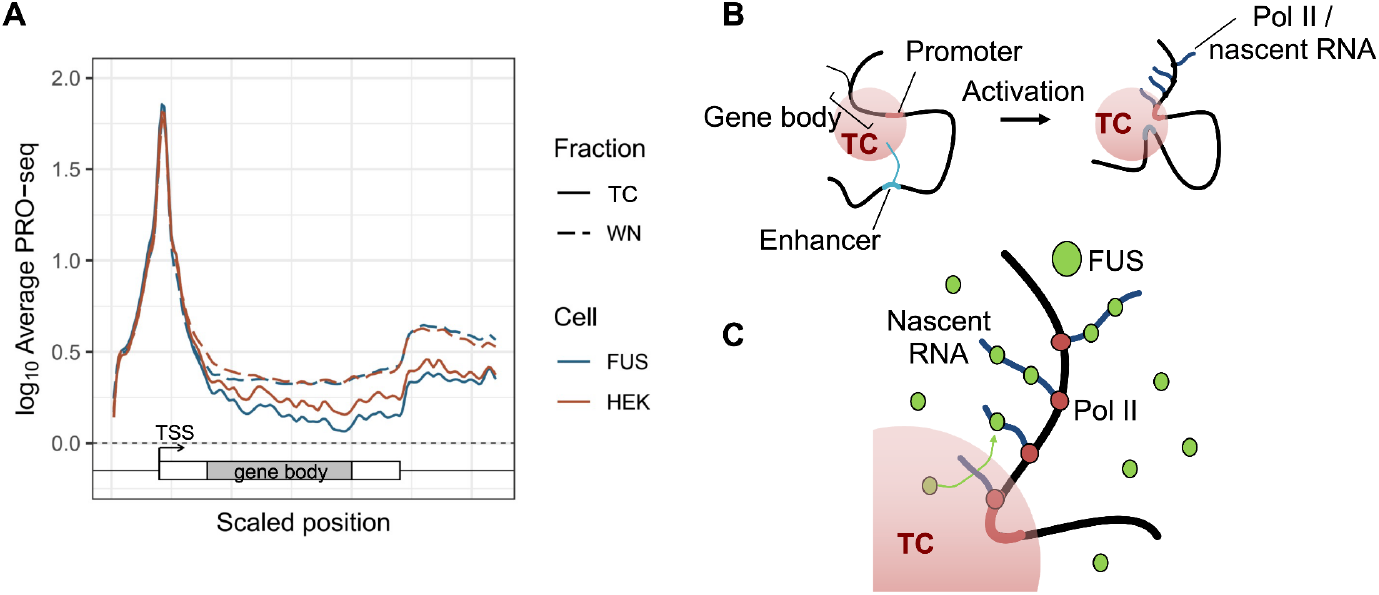
FUS overexpression leads to depletion of the gene body from TC and the models. **A**. Scaled average metagene profile of PRO-seq performed on TC isolates and whole nuclei (WN) in GFP-FUS overexpressed HEK293 (FUS) and wildtype HEK293 (HEK). **B**. Dynamic model of transcriptional elements in and out of TCs upon activation. **C**. FUS facilitated gene body escape.

In summary, our approach of sequencing chromatin in nuclear condensates has provided valuable insights into the behavior of chromatin within these structures. Our results suggest a dynamic model in which promoters and genes reside within TC, and upon activation, enhancers move into TC while the actively transcribing gene body moves out of it (Fig 4B). The process of gene body export is facilitated by the interaction between FUS protein and nascent RNA, which prevents both from condensing into phases (Fig 4C). These findings contribute to a better understanding of the regulation of gene expression through phase-separated biophysical condensates and the role of FUS in the process.

Our model is consistent with existing theories and supports predictions of the phase separation model for transcriptional regulation (Hnisz et al., 2017). We utilized a strategy to isolate nuclease-resistant particles, which we hypothesized to be nuclear condensates (Fig 1). These nuclear condensates were significantly enriched with active histone marks, indicating the presence of activating TCs. While TCs are expected to have higher local concentrations of transcription factors, elongating Pol II-nascent RNA complexes must move to other compartments for processing and mRNA export. Our main finding that gene bodies are generally exported out of TC upon activation supports this prediction (Fig 2), shuttling to elongation and processing condensates that may be much smaller than TCs (Palacio and Taatjes, 2022) and may have evaded detection in our sedimentation fraction.

We propose a novel function for FUS in regulating chromatin dynamics during transcription elongation. FUS is an RNA-binding protein frequently mutated in ALS, and is involved in RNA processing and export (Lagier-Tourenne et al. 2010; Patel et al. 2015; Masuda et al. 2015). FUS contains an intrinsically disordered region (IDR) and can initiate phase condensation by itself, but this process is inhibited when it binds to RNA (Maharana et al. 2018). Our study demonstrated that nascent RNA in the elongating Pol II complex is the primary component that prevents FUS from condensing (Fig 3).

We hypothesized that this prevention of condensation is mutual, and FUS can protect elongating Pol II, leading to its redistribution out of TC. The reduction of gene body nascent RNA in the TC fraction due to FUS overexpression supports this redistribution (Fig 4). Our model also aligns with a previous study that found FUS interacts with the Serine 2 phosphorylated (S2P) C-terminal domain (CTD) of Pol II, where FUS depletion led to abnormal accumulation of S2P Pol II near promoters (Schwartz et al. 2012). Therefore, our model suggests that a lack of FUS may cause S2P Pol II to become trapped within TCs, resulting in accumulation near promoters. Together, these studies offer a more comprehensive understanding of FUS’s role in nascent RNA elongation and highlight the importance of protein-RNA interactions in regulating transcriptional regulation.

## MATERIALS AND METHODS

### Cell lines and culture

HeLa cells were obtained from ATCC. Flp-In-293 cell line and related plasmids was a generous gift from Dr. Andrew Grimson’s laboratory (Cornell University). GFP-FUS plasmid was a generous gift by Dr. Fenghua Hu (Cornell University). Cell lines were cultured in Dulbecco’s Modified Eagle Media (DMEM) supplemented with 10% Fetal Bovine Serum and 1% Penicillin/Streptomycin.

#### Generation of FUS-HEK293 stable cell lines

The coding region of GFP-FUS or GFP was cloned into the multi-cloning site of pcDNA5-FRT vector using the InFusion cloning. system (Takara). Flp-In-293 cells were maintained in DMEM with high glucose and L-glutamine supplemented with 10% Fetal Bovine Serum, 1% Penicillin/Streptomycin, and 100 ug/ml Zeocin. To generate GFP-FUS of GFP stable cell lines, Flp-In-293 cells in 6 well plate were transfected with 1.8 ug pOG44 plasmid, 0.2 ug of pcDNA5/FRT/GFP-FUS or pcDNA5/FRT/GFP plasmid, and 6 ul of FuGENE/HD reagent in 100 ul OPTIMEM media. 48 hours after the transfection, cells were split into 10 cm dishes and selected by adding 200 ug/ml Hygromycin to the growth media. After 1 week, hygromycin resistant colonies were retrieved and cryopreserved.

### Physical isolation of nuclease-resistant nuclear condensate fraction

#### Isolation of nuclei

Approximately 4 million cells were collected from 10 cm culture dishes by rinsing the plates with cold PBS twice and applying 1.5 mL of buffer A (100 mM Tris-HCl pH 7.4, 100 mM NaCl, 30 mM MgCl2, 0.2% Triton, 0.5 mM DTT, 100 mM sucrose) to the plates. The cells were then scraped off and transferred directly to dounce homogenizers, where they were homogenized by applying 40 strokes, 1 second up and 1 second down using the tight pestle (pestle B). After adding 1.5 mL of buffer B (100 mM Tris-HCl pH 7.4, 100 mM NaCl, 30 mM MgCl2, 0.1% Triton, 0.5 mM DTT, 250 mM sucrose) and mixing, the A/B buffer cell pool was layered onto the top of 8 mL of 1 M sucrose cushion in a 15 mL falcon tube, with minimal mixing of the cell suspension with the sucrose cushion. The samples were then centrifuged at 1000 x g for 5 min at 4°C to pellet nuclei, and supernatants were discarded.

#### Light nuclease digestion

The nuclei were resuspended in 250 μL of MNase buffer (50 mM Tris-HCl, 5 mM CaCl2 pH 7.9 @ 25°C) supplemented with 100 μg/mL BSA and the resuspended nuclei were re-pooled. A 50 μL mix was taken out and saved as a whole nuclei (WN) fraction. The remaining 200 μL sample was sonicated for 20 total seconds, with 1 second on and 1 second off at 50% power, resulting in 10 seconds of total sonication. A 10 μL mix was taken out and saved as sonicated untreated nuclei for quality control analysis. To the 200 μL sonicated nuclei, 4 μL of the 1.25 U/μL MNase enzyme dilution in 1x MNase buffer was added. The samples were incubated for 5 minutes at 37°C. The samples were immediately placed on ice and 12.5 μL of 100 mM EGTA was added at a final concentration of 6.25 mM to terminate MNase activity. A 10 μL of the mix was saved for the quality control of sonicated treated nuclei.

#### Nuclear Condensate sedimentation

∼200 μL of light MNase-treated fraction was spun at 3000 x g for 3 minutes. Then, 50 μL of the supernatant from the light MNase-treated fraction should be taken out and saved for quality control. All remaining supernatants were removed. The pellet was resuspended in 50 μL of buffer D (50 mM Tris-Cl pH 8.0, 25% (v/v) glycerol, 5 mM MgAc2, 0.1 mM EDTA, 5 mM DTT) for nuclear condensate (NC) fraction. 50 μL of WN fraction from the previous step was also spun at 3000 x g for 3 minutes and resuspended in 50 μL of buffer D. 5 μL of the NC and WN fractions were saved for quality control analysis. Finally, the WN and NC fractions were rapidly frozen in liquid nitrogen and stored at -80°C.

### Immunoblotting

Anti-Histone 4 pan-acetyl (H4ac) antibody was purchased from ActiveMotif. Anti-Histone 3 (H3) and other antibodies were purchased from Abcam. For immunoblotting, 20 μg, 10 μg, 5 μg, 2.5 μg, 1.25 μg, and 625 ng of protein from WN and NC fractions were denatured in 1x Laemmli buffer at 95°C and electrophoresed in 10% SDS-PAGE in 1x Tris-Glycine buffer. The separated proteins were then transferred to a PVDF membrane and blocked with 5% skim milk in TBS-T buffer (20 mM Tris-HCl, 150 mM NaCl, 0.1% Tween-20, pH 7.5). After washing in TBS-T, the membrane was incubated overnight at 4°C in a 1:5000 dilution of primary antibody in the blocking buffer, followed by 1:10,000 HRP-conjugated secondary antibody. The immunoblots were developed using an enhanced chemiluminescence system.

### DNA-seq

Genomic DNA from the whole nuclei or nuclear condensate fractions were extracted using genomic DNA extraction kit (New England Biolabs). DNA-seq library was prepared from 50 ng of extracted genomic DNA using Illumina Nextera DNA Library Prep kit. DNA-seq experiments were performed in biological replicates.

### Precision Run-On sequencing (PRO-seq)

PRO-seq was performed on whole nuclei or nuclear condensate fractions in the storage buffer (Buffer D) as described previously (Kwat et al, 2013), with the following modifications. Briefly, isolated fractions were incubated in the nuclear run-on reaction condition (5 mM Tris-HCl pH 8.0, 2.5 mM MgCl_2_, 0.5 mM DTT, 150 mM KCl, 0.5% Sarkosyl, 0.4 units / μl of RNase inhibitor) with biotin-NTPs and rNTPs supplied (18.75 μM rATP, 18.75 μM rGTP, 1.875 μM biotin-11-CTP, 1.875 μM biotin-11-UTP for uPRO; 18.75 μM rATP, 18.75 μM rGTP, 18.75 μM rUTP, 0.75 μM CTP, 7.5 μM biotin-11-CTP for pChRO) for 5 min at 37°C. Run-On RNA was extracted using TRIzol, and fragmented under 0.2 N NaOH for 15 min on ice. Fragmented RNA was neutralized, and buffer exchanged by passing through P-30 columns (Biorad). 3′ RNA adaptor (/5Phos/NNNNNNNNGAUCGUCGGACUGUAGAACUCUGAAC/3InvdT/) is ligated at 5 μM concentration for 1 hours at room temperature using T4 RNA ligase (NEB), followed by 2 consecutive streptavidin bead bindings and extractions. Extracted RNA is converted to cDNA using template switch reverse transcription with 1 μM RP1-short RT primer (GTTCAGAGTTCTACAGTCCGA), 3.75 μM RTP-Template Switch Oligo (GCCTTGGCACCCGAGAATTCCArGrGrG), 1x Template Switch Enzyme and Buffer (NEB) at 42°C for 30 min. After a SPRI bead clean-up, the cDNA is PCR amplified up to 20 cycles using primers compatible with Illumina Small RNA sequencing. PRO-seq experiments were performed at least in 3-biological replicates.

### Analysis of chromatin dynamics using DNA-seq and PRO-seq data

#### DNA-seq analysis

DNA-seq results were aligned to the hg38 genome using BWA aligner in paired-end mode. Aligned DNA-seq reads were normalized to fragments per million (FPM) per genomic regions as defined by promoter proximal (−500 bp to 500 bp), gene body (+1 kb from annotated transcription start site to cleavage polyadenylation site), or other regions defined by ChromHMM (see ChromHMM analysis). Boxplots of nuclear condensate (NC) / whole nuclei (WN) fractions were generated from the normalized FPM per region.

#### PRO-seq analysis

PRO-seq results were aligned to the hg38 genome using STAR aligner, and further processed using custom scripts to strand specific genome-wide fragment counts per base data in bedgraph format. PRO-seq reads were normalized to FPM per genomics regions of promoter proximal, gene body or other ChromHMM regions as described in the DNA-seq analysis. Boxplots were generated from normalized FPM per region. Scaled average profiles were produced as described previously (Kwak et al, 2013), using 200 bins in the gene body regions of 500 bp downstream from the TSS to 500 bp upstream of the polyadenylation sites in all expressed genes (gene body FPM > 0.1) longer than 10 kb.

#### ChromHMM analysis

ChromHMM analysis of HEK293 cells were performed as described previously (Ernst and Kellis, 2012) using H3K4me1, H3K4me3, H3K9me3, H3K36me3, and H3K27me3 ChIP-seq data (Encode ENCSR000FCG, ENCSR000DTU, ENCSR000FCJ, ENCSR910LIE, and Chip-Atlas DRX013192, respectively). From the 15 state ChromHMM annotation, we combined TssA and TSSAFlnk as Promoters (Prm); TxFlnk, Tx, and TxWk asTranscribed gene bodies (Txn); EnhG and Enh as Enhancers (Enh); TSSBiv, BivFlnk, and EnhBiv as Bivalent (Biv); ReprPC, and ReprPCWk as Repressed (Rpr). This resulted in a total of 8 states: Promoter (Prm), Transcribed (Txn), Enhancer (Enh), Repetitive (Rpt), Heterochromatin (Het), Bivalent (Biv), Repressed (Rpr), and Quiescent (Qes) regions. Boxplots of chromatin enrichment in nuclear condensates were calculated based on this 8 region annotations.

### Live cell fluorescence microscopy of FUS condensation

Flp-In-293 cells stably expressing GFP-FUS were integrated with a Zeiss 710 incubation system to maintain 37°C and 5% CO2. Cells were treated with Flavopiridol at a final concentration of 500 nM and then immediately imaged for the next 1 hour. Live-cell imaging was conducted using a Zeiss LSM710 confocal microscope equipped with a 488 nm excitation laser and a 63x oil immersion objective lens. The system was integrated with a Zeiss 710 incubation system to maintain 37°C and 5% CO2. Images were acquired at a 600 fps frame rate and 512 × 512-pixel resolution, with 25 z-stacks captured in 6 μm sections. Data analysis was performed using LASX software to quantify GFP-labeled proteins and assess their localization.

## Supporting information

Supplementary Media

## DATA AVAILABILITY

Data is deposited to Gene Expression Omnibus (GEO) under the accession number TBD.

## ACKNOWLEDGEMENTS

We thank Drs. Fenghua Hu and Andrew Grimson (Cornell University) for generously sharing FUS plasmid and reagents for the Flp-In system. We thank the current and the previous members of the Kwak lab and the Department of Molecular Biology and Genetics at Cornell University for providing constructive discussion and sharing unpublished datasets for this study. We also thank Cornell BRC Imaging Facilities for the assistance in live-cell microscopy acquisition. This study was supported by discretionary funds to HK (Cornell University).

## AUTHOR CONTRIBUTIONS

HK conceived the study. SAL, SSYK, and MZ performed experiments. SAL, EGU, and HK performed computational analysis and wrote the manuscript. All authors read and approved the final manuscript.

