## Supplementary Information for "FUS Facilitates Gene Body Export from Nuclear Condensates"

**Figure S1**

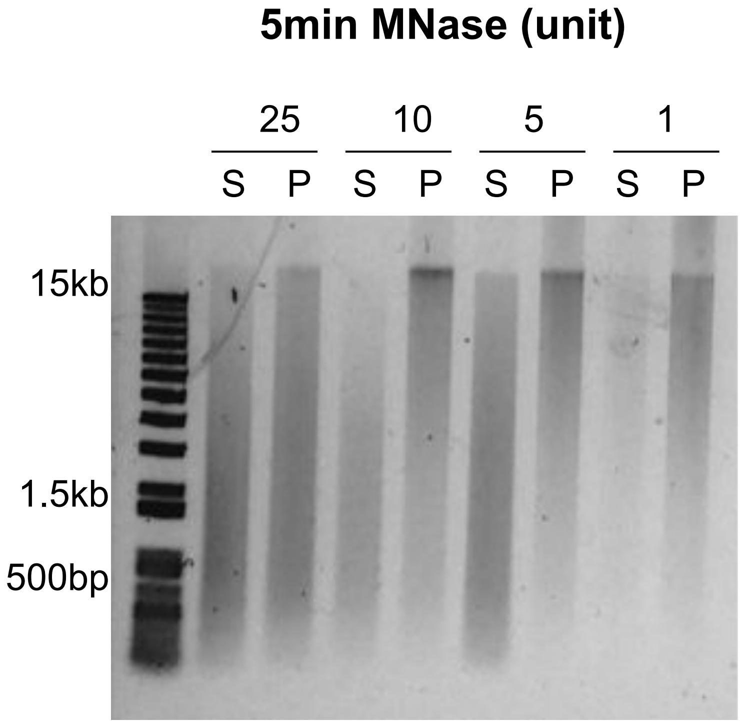


**Fig S1.** Titration of nuclease digestion to differentiate soluble nucleoplasm (S) and pelleted condensate (P) fractions. Agarose gel electrophoresis of the fractionated chromatin shows the sizes of digested DNA in varying units of micrococcal nuclease (MNase).

**Figure S2**

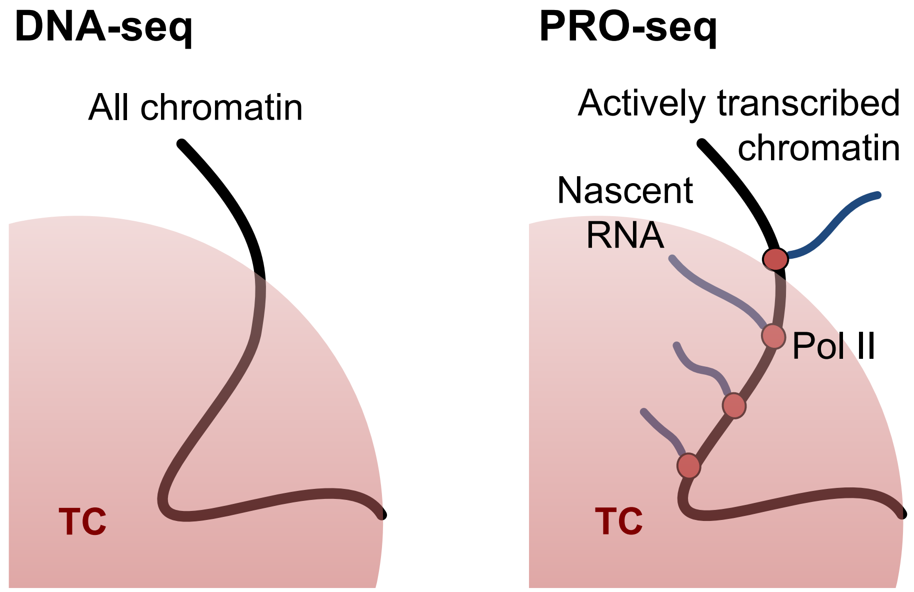


**Fig S2.** Schematics of differentiating actively transcribing chromatin from all chromatin by comparing TC DNA-seq and TC PRO-seq.

**Figure S3**

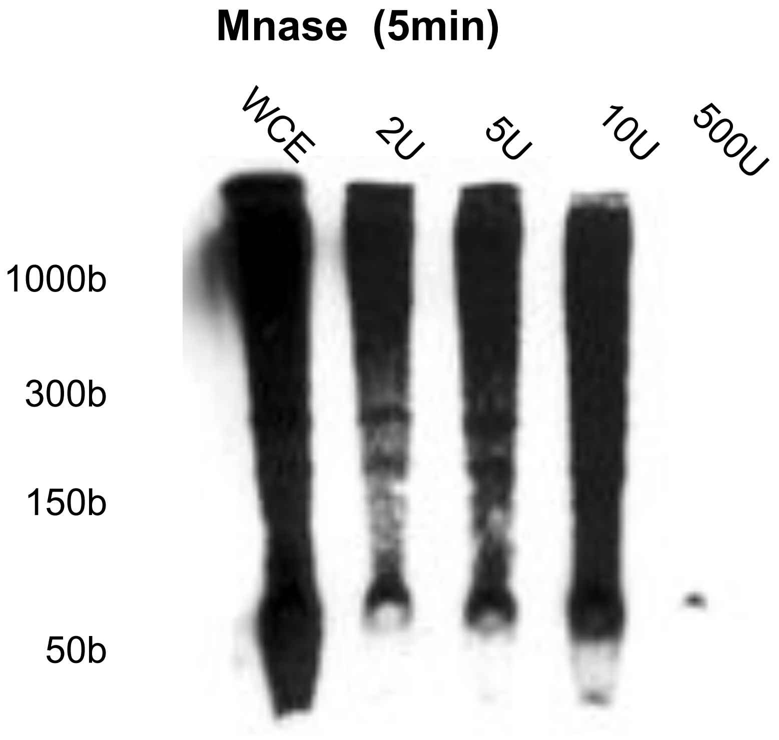


**Fig S3.** Transcriptional Run-on in nuclease digested condensates. Nuclear run-ons using biotin-CTP substrates were performed on pelletted condensate fractions with varying amounts of MNase treatment. The biotin run-on RNA were transferred to the nylon membrane after electrophoresis and blotted with streptavidin-HRP substrate. Whole cell extract (WCE) was used as a positive control.

### Figure S4

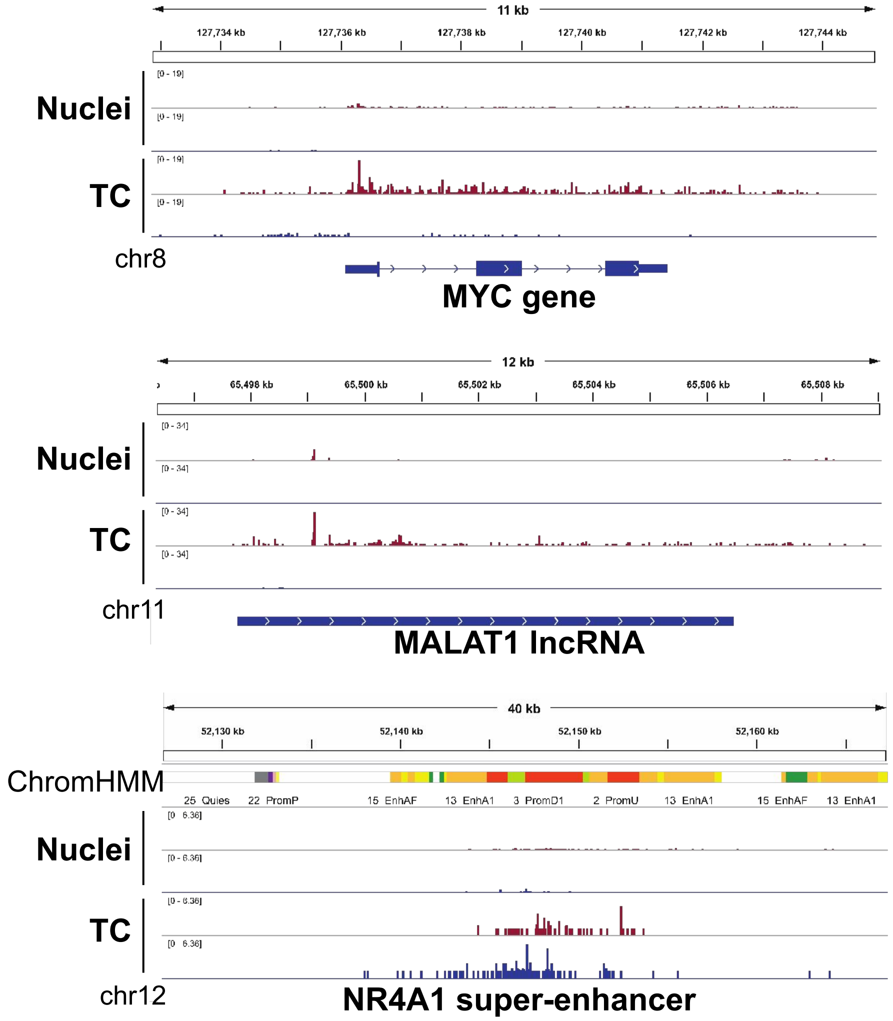


**Fig S4.** Genes and non-coding RNA loci enriched in TCs. Example genome browser views of whole nuclei and TC PRO-seq at a gene, lncRNA, and a super-enhancer. The y-axis readcount scales were normalized to the total readcount of the library.

**Figure S5**

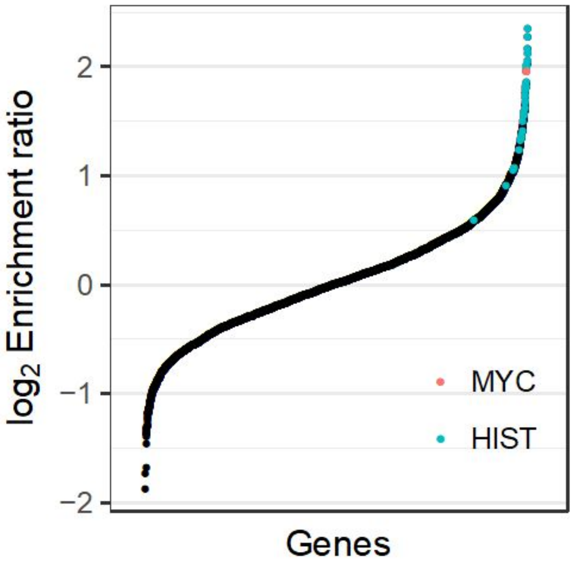


**Fig S5.** Enrichment of MYC and histone genes(HIST) in TC PRO-seq. Y-axis: log2 enrichment ratio of TC / whole nuclei.

**Figure S6**

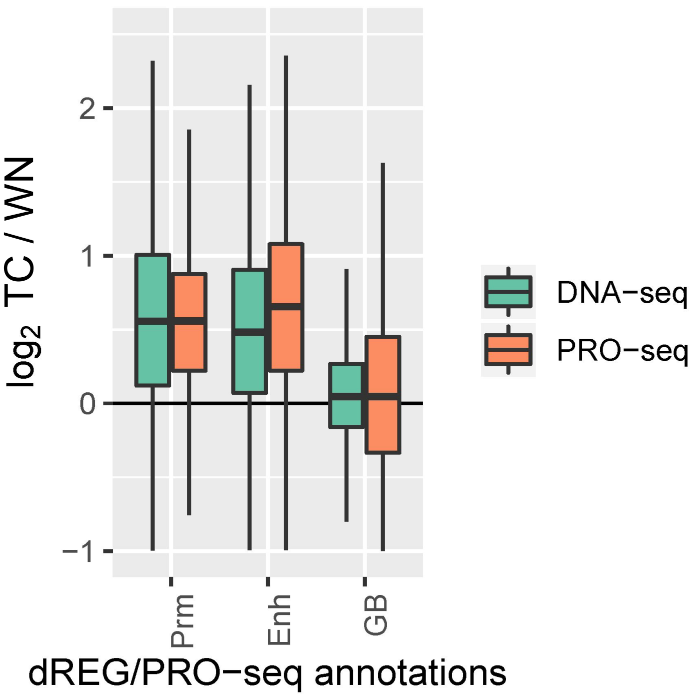


**Fig S6.** Enrichment of dREG annotated promoters, enhancers, and gene bodies in TC PRO-seq data. Data was normalized with respect to all transcribed regions, mostly gene bodies (GB), therefore gene bodies appeared relatively unchanged in this normalization.

**Figure S7**

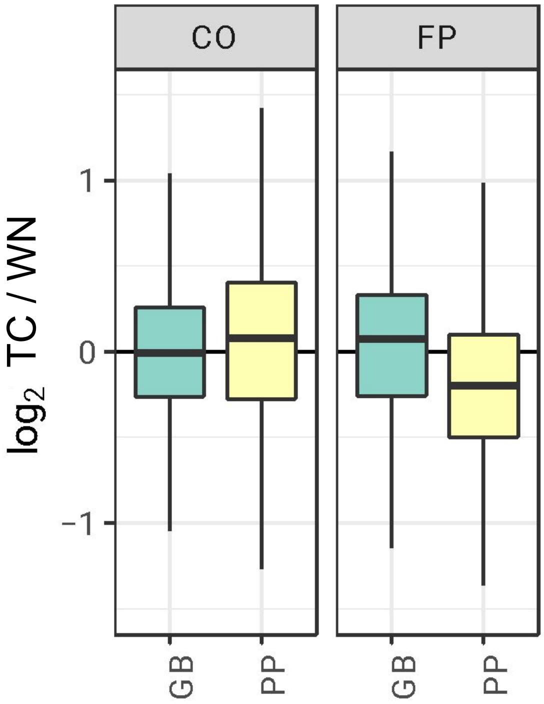


**Fig S7.** Enrichment of gene body (GB) and promoter proximal (PP) regions in TC without (CO) and with flavopiridol treatment (FP). PRO-seq data was normalized with respect to all transcribed regions, mostly gene bodies (GB), therefore gene bodies appeared relatively unchanged in this normalization.

#### Supplemental Media

**Supplemental Media.** Video of FUS condensation upon FP treatment. Real time counter of minutes:seconds is located at the upper left corner.
